# AI^2^BMD: efficient characterization of protein dynamics with *ab initio* accuracy

**DOI:** 10.1101/2023.07.12.548519

**Authors:** Tong Wang, Xinheng He, Mingyu Li, Yusong Wang, Zun Wang, Shaoning Li, Bin Shao, Tie-Yan Liu

**Affiliations:** Microsoft Research AI4Science, Beijing, 100080, China; Work done during an internship at Microsoft Research; State Key Laboratory of Drug Research and CAS Key Laboratory of Receptor Research and, Shanghai Institute of Materia Medica, Chinese Academy of Sciences, Shanghai, 201203, China; Department of Pathophysiology, Key Laboratory of Cell Differentiation and Apoptosis of Chinese Ministry of Education, Shanghai Jiao Tong University, School of Medicine, Shanghai, 200025, China; Institute of Artificial Intelligence and Robotics, Xi’an Jiaotong University, Xi’an, 710049, China

## Abstract

Biomolecular dynamics simulation is a fundamental technology for life sciences research, and its usefulness depends on its accuracy and efficiency. Classical molecular dynamics simulation is fast but lacks chemical accuracy. Quantum chemistry methods like density functional theory (DFT) can reach chemical accuracy but cannot scale to support large biomolecules. We introduce an <u>AI</u>-based *<u>a</u>b initio* <u>b</u>io<u>m</u>olecular <u>d</u>ynamics system (AI^2^BMD) that can efficiently simulate large biomolecules with *ab initio* accuracy. AI^2^BMD uses a protein fragmentation scheme and machine learning force field to achieve generalizable *ab initio* accuracy for energy and force calculations for various proteins comprising over 10,000 atoms. Compared to DFT, it reduces computational time by several orders of magnitude. With several hundred nanoseconds of dynamics simulations, AI^2^BMD demonstrated its capability of efficiently exploring the conformational space of peptides and proteins, deriving accurate ^*3*^*J-couplings* that match NMR experiments, and showing protein folding and unfolding tendencies. Furthermore, AI^2^BMD enables precise free energy calculations for protein folding, and the estimated melting temperatures are well aligned with experiments. AI^2^BMD could potentially complement wet-lab experiments, detect the dynamic processes of bioactivities, and enable biomedical research that is currently impossible to conduct.

## MAIN

The paradigm of life sciences research is shifting as the accuracy of the computational simulation models is getting indistinguishable from that of wet-lab experiments^1,2^. Among the computational models, as the “computational microscope”, molecular dynamics simulation is of particular interest for understanding how life works^3-5^. Molecular dynamics simulations study the dynamic evolution of molecules by moving the atoms, or more specifically, the nuclei in a molecular system. They differ in the way the forces are calculated^5^. In classical molecular dynamics (termed as “MD”), forces are calculated using a prescribed interatomic potential function, while in *ab initio* molecular dynamics (termed as “AIMD”), forces are calculated using the potential derived from the electronic structure of the molecules^6^. Although AIMD provides accurate characterization of molecules, the main challenge of applying AIMD to simulate biomolecules is scalability. On the one hand, the widely used quantum chemistry methods for AIMD are very computationally expensive, e.g., the time complexity of density functional theory (DFT) is about *O*(*N*^3^) and that of CCSD(T) is even *O*(*N*^7^). On the other hand, to observe significant conformational changes for biomolecules such as proteins, it usually takes billions of steps with at least cubic time complexity for thousands of atoms^7^. Till now, scalable and accurate AIMD for biomolecules does not exist.

To alleviate the dilemma, in recent years, machine learning force fields learning from data generated at DFT level bring accurate calculation with much less cost and are applied for small peptides and proteins^8-10^. However, generalization ability is the key challenge of the applicability and robustness for biomolecule simulations^11^. First, as the conformational space of a molecule is enormous, training on limited conformations and adapting it for conformational space exploration is difficult^4^. Second, since training data generated by DFT is cubically increased with the molecular sizes, lack of training data hinders MLFFs’ application for large biomolecules^11^. Furthermore, it is impossible to train a specific model for each kind of protein, and thus a unified solution with good generalization to various proteins is needed.

In this study, we propose AI^2^BMD, a generalizable solution to efficiently simulate various proteins with *ab initio* accuracy (Figure 1). A generalizable protein fragmentation approach splits proteins into overlapped protein units. Simulations are performed by the AI^2^BMD simulation system while in each simulation step, the AI^2^BMD potential based on ViSNet^9^ calculates the energy and atomic forces for the protein with *ab initio* accuracy. By comprehensive analysis from both kinetics and thermodynamics, AI^2^BMD exhibits good alignments with wet-lab experiment data such as the melting temperature of fast folding proteins and different phenomenon compared with molecular mechanics.

**Figure 1.**
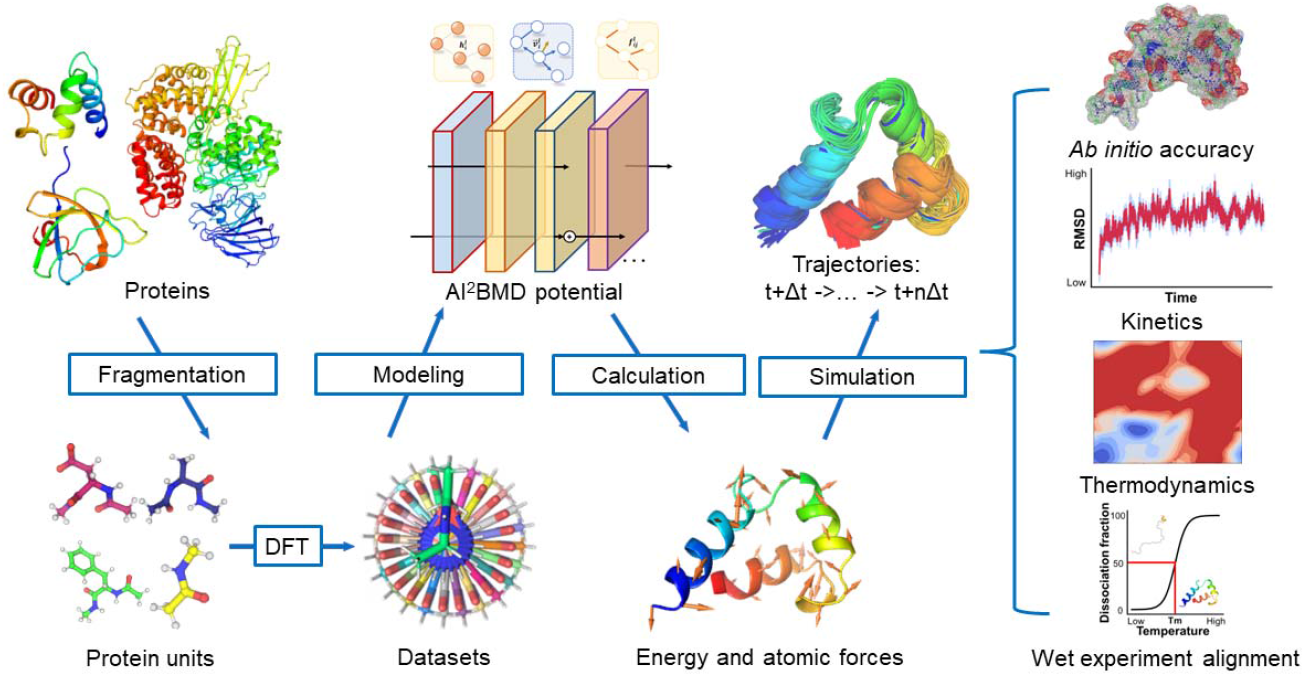
The overall pipeline of AI^2^BMD. Proteins are divided into protein units by fragmentation process. The AI^2^BMD potential is designed based on ViSNet, and the datasets are generated at DFT level. It calculates the energy and atomic forces for the whole protein. The AI^2^BMD simulation system is built upon all these components and provides a generalizable solution to perform simulations for various proteins. It makes *ab initio* accuracy on energy and force calculations. By comprehensive analysis from both kinetics and thermodynamics, AI^2^BMD exhibits good alignments with wet-lab experiment data and detects different phenomenon compared with molecular mechanics.

### Accurate and efficient energy and force calculations

As a generalizable solution to accurately simulate various proteins, AI^2^BMD first adopted a generalizable protein fragmentation approach. As demonstrated above, directly generating training data at DFT level for whole large proteins is computational prohibitive. Thus, we fragmented proteins into protein units (i.e., dipeptides), calculated intra- and inter-unit interactions, and then reassembled them to determine protein energy and force of the conformation (see Methods for more details). On the one hand, the set of protein units can form all kinds of proteins, which indicates it is a generalizable fragmentation approach. On the other hand, our fragmentation approach contains only 21 kinds of protein units, and all protein units have similar and moderate numbers of atoms (range from 12 to 36), which is convenient for DFT data generation and machine learning force field training. We then built a comprehensively sampled protein unit dataset. During dataset construction, we meticulously scanned the mainchain dihedrals of all protein units to cover a wide range of conformations and run AIMD simulations at M06-2X/6-31g* level^12^ since the functional models dispersion well and has been widely used in previous studies for biomolecules, thereby obtaining 20.88 million samples with *ab initio* accuracy (see Methods for more details). The whole dataset was split into training, validation and test sets to train ViSNet^9^ models as AI^2^BMD potential. The model is designed to fully encode physics-informed molecular interactions by elaborate model operations and calculates 4-body interactions with linear time complexity, which subsequently generates precise force and energy estimations based on the atom types and the coordinates as inputs (Figure S1 and Methods).

The performance of AI^2^BMD potential was compared with that of the conventional molecular mechanics (MM) force field on the test set, with the results presented in Table S1. In terms of energy mean absolute error (MAE), AI^2^BMD potential outperformed the MM force field by approximately two orders of magnitude (AI^2^BMD: 0.023 ± 0.010 kcal/mol, MM: 3.198 ± 1.091 kcal/mol). Similarly, AI^2^BMD potential also demonstrated superior performance of the force MAE (0.036 ± 0.004 kcal/mol*Å^-1^) compared to MM (8.125 ± 0.658 kcal/mol*Å^-1^). Overall, AI^2^BMD potential offers accurate predictions for both potential energy and atomic forces for protein units.

Based on AI^2^BMD potential, we developed an MD simulation system and conducted simulations in an explicit solvent for nine proteins with the number of atoms ranging from 175 to 13,728 (Figure 2a and more details in Method). Each protein was assessed with 5 folded, 5 unfolded, and 10 intermediate structures derived from replica exchange MD simulations as the initial conformations, and 10 AI^2^BMD simulation steps were run resulting in 200 structures per protein. The AI^2^BMD simulation system’s capability was evaluated by comparing its results to those calculated by the MM force field, with those calculated by DFT as the ground truth values (Figure 2b-2e). For evaluation on potential energy (Figure 2b, 2c), MM exhibited a broader error distribution and a significantly higher upper bound of error (i.e., the max error) compared with AI^2^BMD. The average MAE of MM potential energy consistently hovered around 0.2 kcal/mol per atom, whereas AI^2^BMD achieved a significantly lower value (0.038 kcal/mol per atom, averaged over the five proteins, Figure 2b). As protein size increased from Chignolin (175 atoms) to PACSIN 3 (1,040 atoms), the increase of energy errors could be attributed to insufficient modeling for the escalating many-body interactions among protein units. For proteins from SS0941 with 2,450 atoms to Aminopeptidase N with 13,728 atoms, the ground truth could only be determined through fragmented DFT (Figure 2c). For these four proteins, AI^2^BMD’s performance (the MAE of 7.18 × 10^−3^ kcal/mol per atom) was significantly superior to that of MM (0.214 kcal/mol per atom). In terms of force (Figure 2d,2e), compared with MM force field, AI^2^BMD significantly aligned more closely with DFT results. For the first five proteins directly calculated by DFT, AI^2^BMD had an average MAE of 1.974 kcal/mol*Å^-1^ compared to MM’s 8.094 kcal/mol*Å^-1^ (Figure 2d). For the last four large proteins, AI^2^BMD achieved an average MAE of 1.056 kcal/mol*Å^-1^, while MM’s value was 8.392 kcal/mol*Å^-1^ across four systems (Figure 2e). We further compared the performance of AI^2^BMD for different conformations. As shown in Figure S2-S4, the MAE of potential energy for unfolded, intermediate and folded conformations of each kind of protein were analyzed, respectively. The MAE of potential energies of different conformations fluctuated among different proteins while those of atomic forces were slightly increased from unfolded conformations to folded conformations.

**Figure 2.**
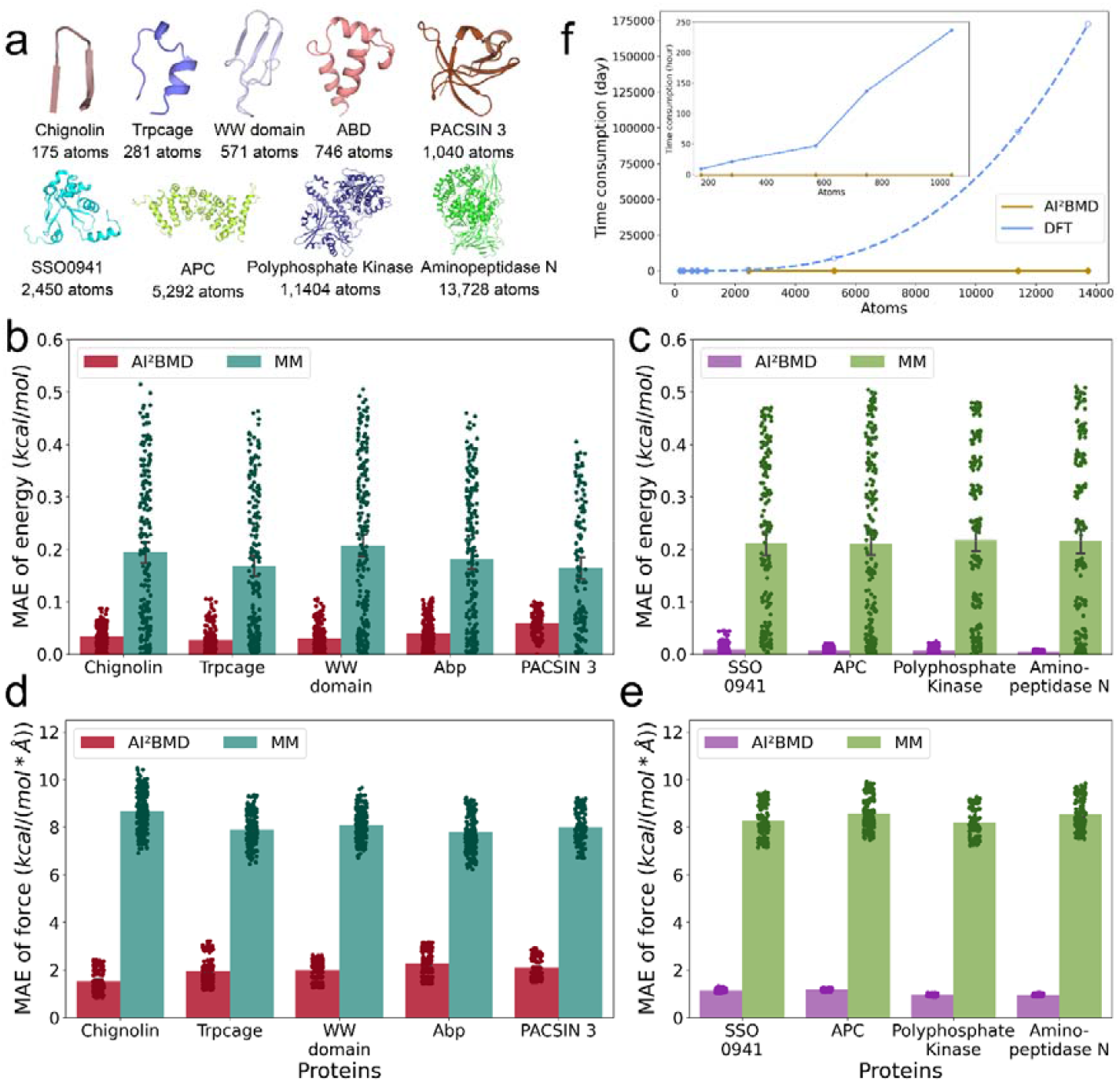
Evaluation of energy and force calculations by AI^2^BMD and molecular mechanics (MM) for various proteins. a, The folded structures of 9 evaluated proteins. For these proteins, the number of atoms ranges from 175 to 13,728. b-e, The mean absolute error (MAE) of potential energy (b-c) and atomic forces (d-e). For each protein, replica exchange MD and structure clustering were performed to select representative structures, e.g., folded, unfolded or intermediate states. Then AI^2^BMD simulations were performed for all representative structures, and 200 samples in total were selected for evaluation. For the first five proteins within 1,040 atoms shown in (b) and (d), DFT calculation for the whole protein is set as the ground truth, while for the last four proteins shown in (c) and (e), the ground truth is set as fragment DFT due to prohibitive computational cost. The MAE of each protein is shown in bar plot while the errors of 200 samples of the protein are shown in scatters. In (b) and (c), the potential energy of each structure subtracts that of a reference structure and then is normalized by the number of atoms. f, Time consumption of energy calculation for 9 proteins. For the last four proteins, the time consumption by DFT was estimated by the fitting curve from those of the first five proteins and was shown in dash line and circles. A subplot in the upper left region exhibits a comparison for the first five proteins.

Furthermore, to examine the efficiency of AI^2^BMD, we compared the time consumption of energy calculation for all 9 proteins. As illustrated in Figure 2f, we present the average computation time for AI^2^BMD and DFT on a single Intel CPU core@2.60GHz with 8 threads. It is obvious that our AI^2^BMD achieved *ab initio* accuracy significantly faster than DFT. The computational time for AI^2^BMD increased nearly linearly. For instance, a protein with 1,040 atoms required an average of 44.39 ± 10.88 seconds, while the corresponding DFT calculations took over 230 hours on average; although a large gap exists compared with classical MD, AI^2^BMD reduces computational time by four orders of magnitude compared with DFT. In summary, AI^2^BMD is versatile, generalizable to various proteins and offers both *ab initio* accuracy and highly efficient calculation for molecular dynamics simulation.

### Conformational space exploration and protein dynamics

To demonstrate the capabilities of AI^2^BMD for conformational space exploration and protein kinetics, we performed AI^2^BMD simulations for both protein dipeptides and proteins. For each protein unit, we conducted 40 independent AI^2^BMD simulations to sample its conformational space. To promote simulation efficiency, 20 initial structures were first derived from comprehensively sampled MM trajectories. Then each initial structure underwent two AI^2^BMD simulation runs for 1,000 ps, respectively. In total, 720 ns AI^2^BMD simulations were performed for protein units. We first evaluated the accuracy of potential energy and atomic force calculations during the simulations. 125 snapshots were evenly picked from a simulation trajectory and the energy and force were calculated by DFT and MM as the ground truth and for comparison, respectively. Throughout the simulations regardless of kinds of protein units, the relative energy and force of AI^2^BMD and the ground truth calculated by DFT exhibited a high degree of overlap, whereas MM deviated from DFT a lot (Figure 3a-3d, Figures S5). In specific, for the negatively charged protein unit Ace-Glu-Nme (Figure 3a), AI^2^BMD exhibited slight differences compared with DFT during the last 300 ps of the simulation, while MM presented a global difference of around 20 kcal/mol. Furthermore, AI^2^BMD consistently outperformed MM by a large margin for positively charged Ace-Arg-Nme (Figure 3b). In addition, for Ace-Phe-Nme with a benzene ring in the sidechain, the energy calculated by MM displayed high fluctuations and was noticeably different from DFT (Figure 3c). With smaller sidechains, such as Ace-Ser-Nme (Figure 3d), the discrepancy between AI^2^BMD and DFT further diminished, while the gap between MM and DFT remained large. For evaluations on atomic forces, AI^2^BMD also demonstrated significantly higher fidelity to the ground truth values compared to MM for all cases in the simulations (Figures S5). Consequently, AI^2^BMD maintained its accuracy across a diverse range of protein units during simulations.

**Figure 3.**
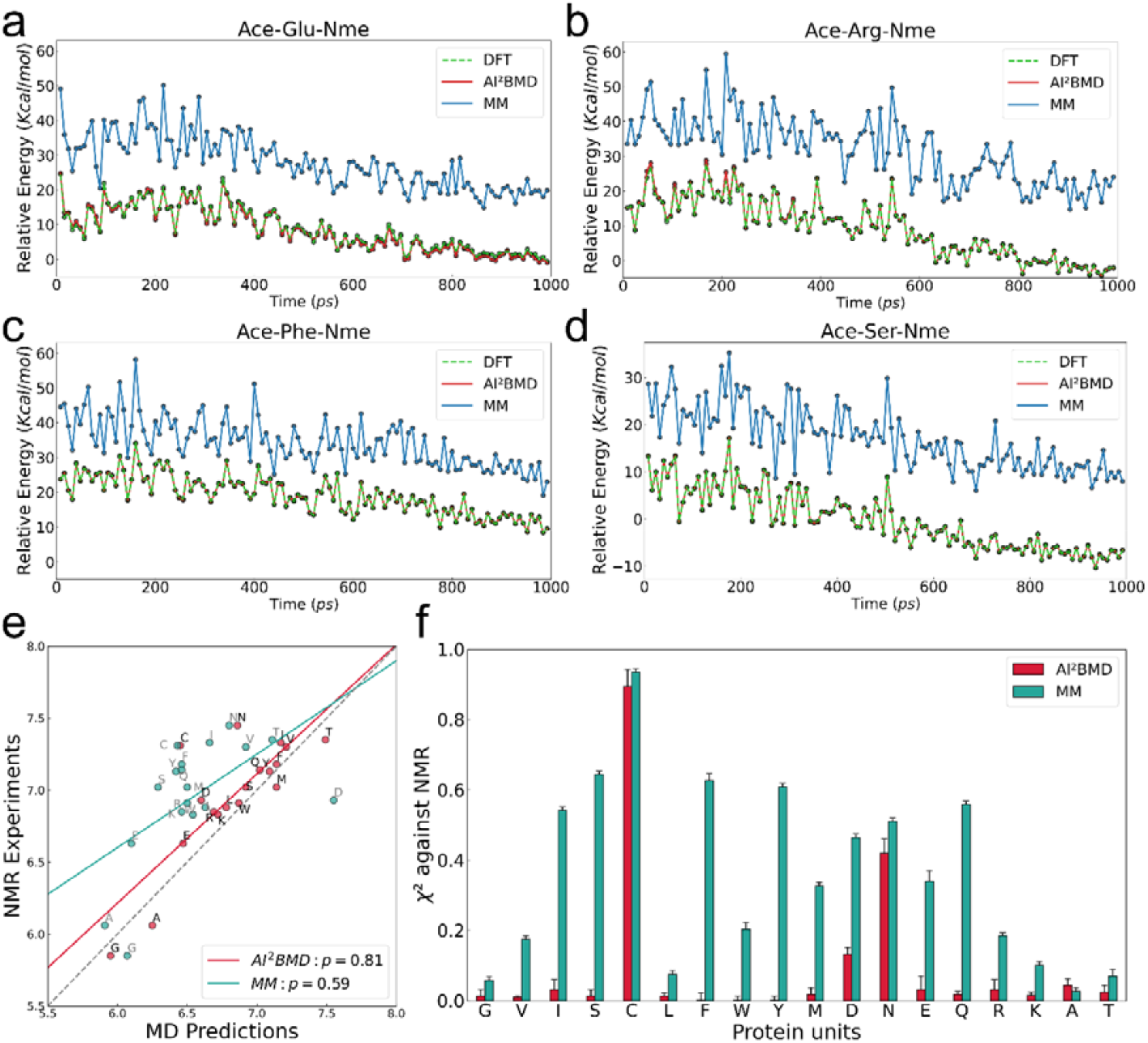
AI^2^BMD simulations for protein units and comparisons with DFT and NMR experiments. a-d, potential energies during the simulations for aromatic Ace-Phe-Nme (a), polar Ace-Ser-Nme (b), negatively charged Ace-Glu-Nme (c) and positively charged Ace-Arg-Nme (d). Calculations by DFT act as the ground truth, and those calculated by MM are shown for comparison. The relative energy of each structure is then calculated by subtracting the energy of the initial structure. (e) Comparison between NMR ^3^*J*(H_N_,H_**α**_) couplings and those derived from simulations driven by AI^2^BMD and MM. The ^3^*J*(H_N_,H_**α**_) couplings derived from AI^2^BMD and MM are shown in red and green points, respectively. For each approach, a linear regression curve is drawn, and the corresponding Pearson correlation is shown in the legend. (f) χ^2^ errors in reproducing ^3^*J*(H_N_,H_**α**_) couplings for each protein unit measured by NMR. The results made by AI^2^BMD and MM are shown in red and green, respectively.

We further analyzed the explored conformational space by AI^2^BMD via reproducing the ^3^*J*(H_N_, H) coupling measured by Nuclear Magnetic Resonance (NMR) experiments^12,13^. The ^3^*J*(H_N_, H_α_) coupling can accurately reflect the φ angle distributions for peptides^12^ and thus was adopted from an experiment view to measure the conformational space exploration by simulations for the protein units. Except for the proline and histidine dipeptides, we calculated the ^3^*J*(H_N_, H_α_) coupling values for the other 18 kinds of protein units based on the main-chain φ angles derived from AI^2^BMD simulation trajectories and then averaged the values of two parallel repeats to obtain the final estimates. As a comparison, those made by MM were reported in ff19SB force field^14^. Figure 3e illustrates that simulations driven by AI^2^BMD exhibited a higher Pearson correlation (p = 0.81) with NMR experiment results than those made by MM (p = 0.59). Furthermore, AI^2^BMD significantly outperformed MM for most protein units as shown in Figure 3f. AI^2^BMD performed suboptimal for Ace-Cys-Nme, Ace-Asp-Nme and Ace-Asn-Nme compared to other cases, which may be attributed to their unique sidechain properties. For Ace-Ala-Nme, although AI^2^BMD slightly underperformed MM, the calculations by both methods were quite accurate. As a contrast, for other cases, particularly the aromatic protein units including Ace-Phe-Nme, Ace-Trp-Nme and Ace-Tyr-Nme, AI^2^BMD significantly reduced the discrepancy between simulations and experiments. Such results further highlighted the effectiveness of AI^2^BMD for conformation exploration and sampling from a wet-experimental perspective.

We then performed AI^2^BMD simulations for the protein Chignolin^15^ to study the differences of protein dynamics sampled by AI^2^BMD. 1,000 initial structures were selected from a 106 µs well-sampled MD trajectory to represent different conformations^7^, and then 100 ps AI^2^BMD simulations in solvent were performed for each, accumulating a total of 100 ns of simulation trajectories. AI^2^BMD captured both the folding and unfolding tendencies of Chignolin during the simulations. In Figure 4a, AI^2^BMD captured a conformational change from a completely unfolded structure to a curled intermediate state. During this process, the average energy difference between AI^2^BMD and DFT was 0.025 kcal/mol per atom, while MM had an error of 0.132 kcal/mol per atom. In contrast, Figure 4b depicted another transition process with an unfolding tendency from a fully folded hairpin structure with well packed β strands to a near hairpin loop conformation. During this process, AI^2^BMD differed from the ground truth by 0.033 kcal/mol per atom, whereas MM exhibited a deviation of 0.124 kcal/mol per atom. In both simulation processes, compared with MM, AI^2^BMD also exhibited much closer force calculation to DFT as shown in Figure S6.

**Figure 4.**
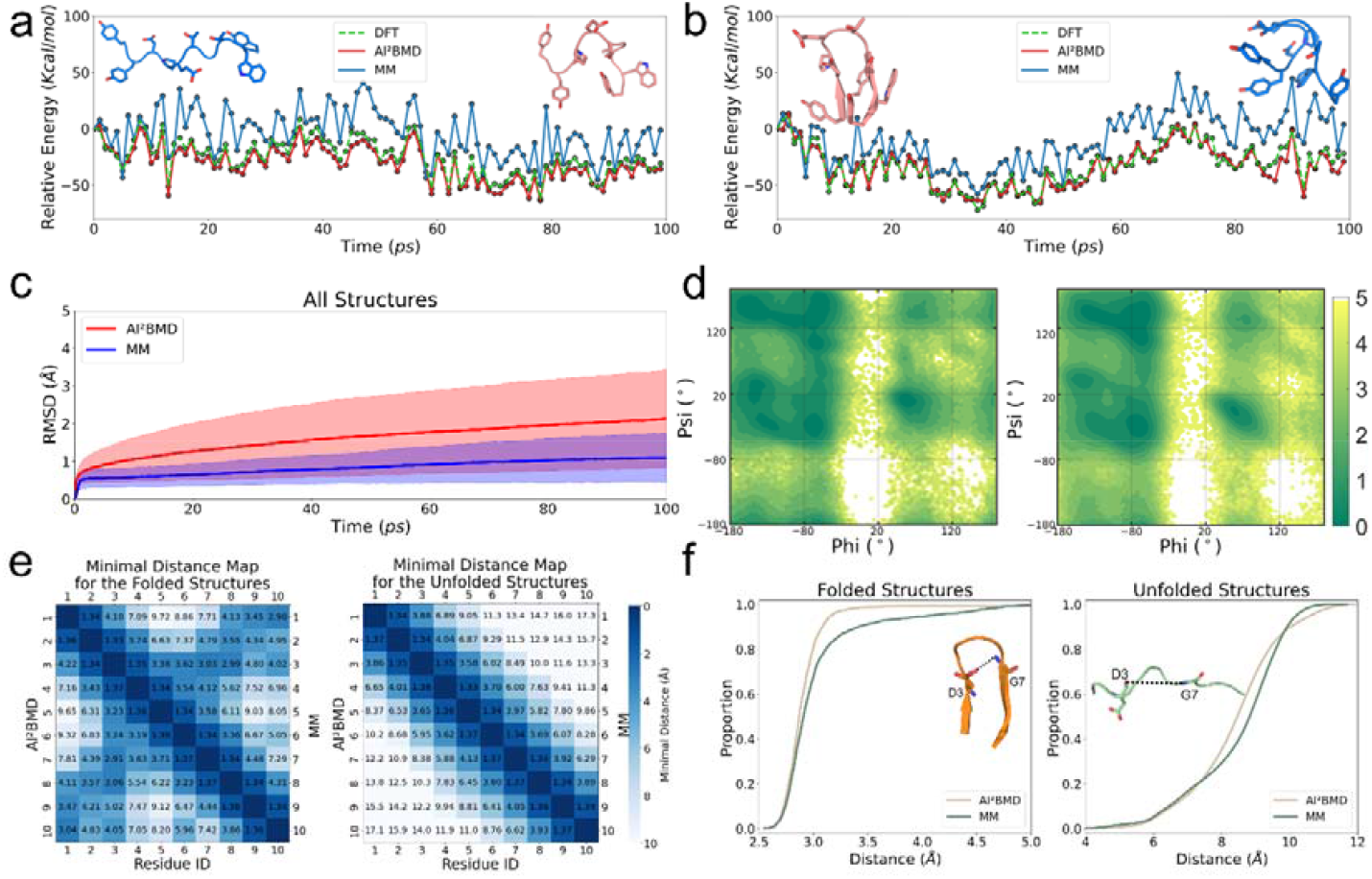
Analysis of protein dynamics of Chignolin by AI^2^BMD simulations. a-b, potential energies during the simulations for Chignolin with two different initial structures. Calculations by DFT act as the ground truth, and those calculated by MM are shown for comparison. The relative potential energy of each structure is calculated by subtracting the energy of the first structure in the simulation. The initial and final structures during the simulations are shown at the top of the panels. c, RMSD during simulations. The average RMSD is shown in line while the standard deviation of all simulation trajectories is shown in shadow. d, Ramachandran plot of conformations sampled by AI^2^BMD (left) and MM (right). e, Residue minimum distance map for folded and unfolded structures. AI^2^BMD (left) and MM (right) are shown in the lower triangle matrix and upper triangle matrix, respectively. f, The cumulative plots for the distance between the main-chain O of D3 and the main-chain N of G7 for folded (left) and unfolded (right) structures, respectively.

To further explore the differences of protein dynamics driven by AI^2^BMD and MD respectively, we also performed simulations by MM with the same initial structures and simulation configurations, and found that AI^2^BMD simulations exhibited several distinct features. First, protein structures fluctuated more in AI^2^BMD, especially over longer simulation times. As shown in Figure 4c, the root mean squared deviation (RMSD) compared to the initial structure increased more rapidly than that in classical MD simulations. After the simulation, AI^2^BMD had an average RMSD of 2.14 Å, while MM simulations reached 1.09 Å. Second, regarding the specific main-chain conformations depicted in the Ramachandran plot (Figure 4d), AI^2^BMD also exhibited a broader distribution than MM, particularly in the domain where phi > 135° and psi ranges from -80° to 180°. This domain corresponds to more flattened protein conformations. The larger RMSD during simulations and the broader Ramachandran plot distribution implies that proteins were more flexibly changed in the machine learning force field with DFT accuracy. Furthermore, using the Q score, which evaluates protein folding based on contacts during simulations^16^, we separated the AI^2^BMD and MM snapshots into folded and unfolded structures. As shown in the residue minimum distance map (Figure 4e), the results of AI^2^BMD and MM were similar in both folded and unfolded structures. Both methods described the stable interactions between Chignolin’s N-terminal and C-terminal aromatic residues and the hydrogen bonds between D3-G7 and D3-T8. For the strongest hydrogen bond between D3 and G7^15,17^, the folded conformations sampled by AI^2^BMD depicted it more stable (the minimum distance of 2.91 Å in AI^2^BMD versus 3.03 Å in MM). The cumulative plots for the distance between the main-chain O of D3 and the main-chain N of G7 (Figure 4f) also illustrate that AI^2^BMD stabilized the hydrogen bond more than MM. For unfolded structures, the trends of AI^2^BMD and MM were similar that both described a much longer distance than that in the folded structures. These results indicate that by performing protein simulations with *ab initio* accuracy, AI^2^BMD can detect both meaningful conformational changes and detailed interatomic interactions to study protein dynamics.

### Free energy and melting temperature estimation

We further investigated the thermodynamic properties of various fast-folding proteins conducted by AI^2^BMD. From the comprehensively sampled trajectories of proteins by D. E. Shaw Research^7^, we evenly procured 100,000 snapshots for each protein and employed the Q score to categorize them into folded and unfolded states. The representative folded and unfolded structures of the seven proteins, i.e., BBA, WW domain, NTL9, homeodomain, protein G, α3D, and λ-repressor are shown in Figure 5a. These proteins consist of 504 to 1,258 atoms and showcase a variety of secondary structures. We first calculated the potential energy values by AI^2^BMD and displayed the potential energy surface using time-lagged independent components (tICs). As shown in Figure 5b for the case of NTL9, for both the results made by MM and AI^2^BMD, folded and unfolded structures were clearly separated with distinct potential energies, while the difference of potential energy between them decreased in the AI^2^BMD’s result. By reweighting on potential energy values, we estimated the free energy difference during protein folding (ΔG) and the melting temperature (Tm) derived from simulation trajectories (Figure 5c, 5d, Table S2). Since such protein simulations were conducted near the corresponding melting temperatures^7^, ΔG is expected to be closer to 0 and calculated Tm is expected to be closer to the simulation temperature. Both MM and AI^2^BMD achieved small free energy differences and comparable calculated melting temperatures to simulation temperatures, while AI^2^BMD slightly outperformed MM for these cases.

**Figure 5.**
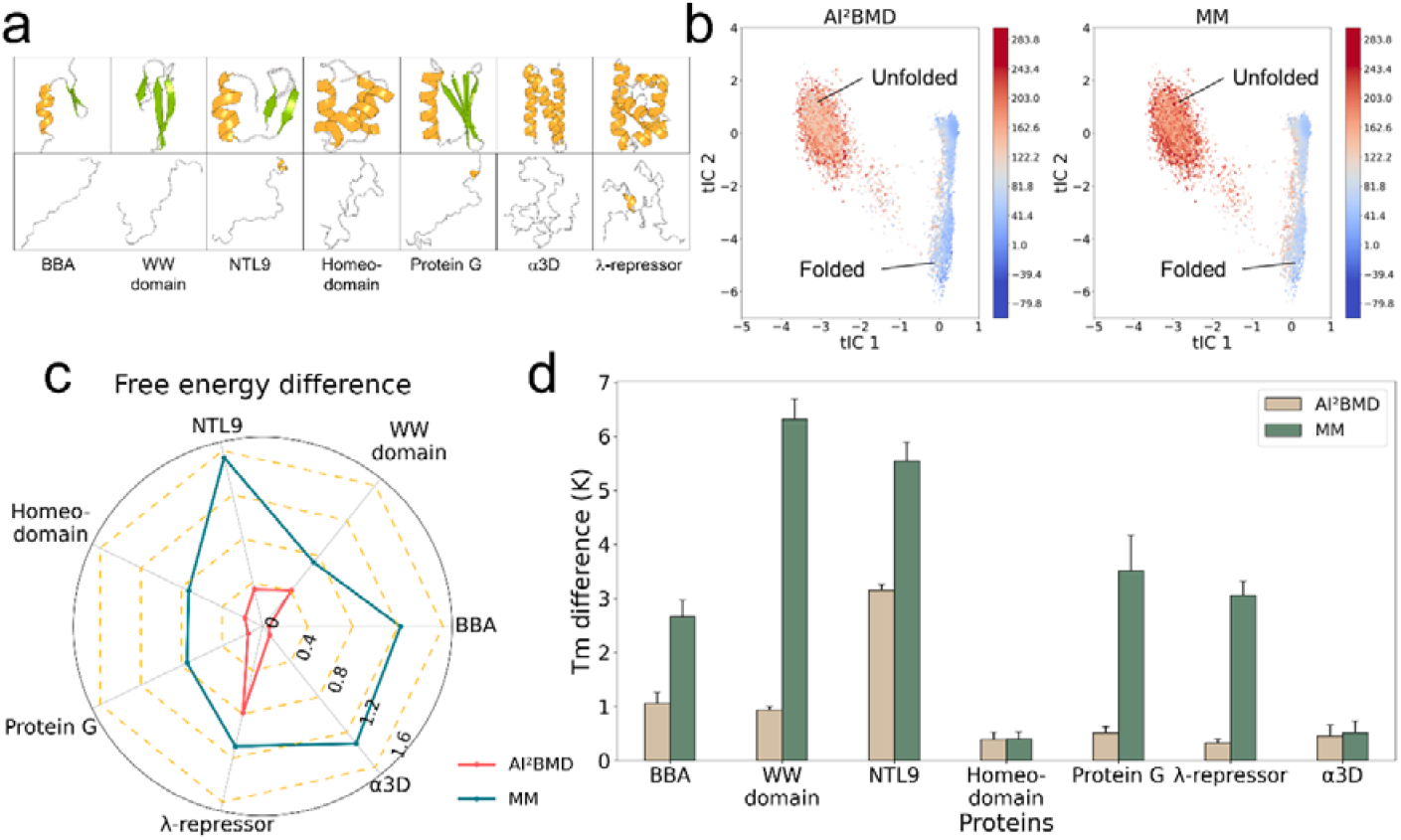
Analysis of free energy calculation and melting temperature estimation for fast folding proteins. a, the representative folded (top line) and unfolded structures (bottom line) for proteins. b, potential energy surface of NTL9 made by AI2BMD (left) and MM (right), respectively. c, the absolute free energy differences between the folded and unfolded structures for proteins. d, The difference between the simulation temperature and the melting temperature (Tm) calculated from simulations.

Notably, WW domain is the only protein dominated by β-sheets in the evaluation^18^. Compared with the experimentally determined melting temperature of 371 ± 2 K^19^, AI^2^BMD provided closer Tm estimation (359.06 ± 0.07 K) compared to MM (353.69 ± 0.38 K). NTL9 has a folded structure with a α-helix and β-sheets^20^. With the simulation temperature of 355 K nearly same to the experimental Tm of 354.75 ± 1.7 K^21^, AI^2^BMD achieved a better estimation of ΔG than MM (−0.34 kcal/mol versus -1.54 kcal/mol). Furthermore, AI^2^BMD estimated the Tm of 351.84 ± 0.11 K, which is more accurate than the value of 349.47 ± 0.35 K made by MM.

In addition, for all alpha proteins, Homeodomain features a folded structure with three α-helices and loops^22^. While AI^2^BMD’s ΔG estimation (−0.18 kcal/mol) is smaller than MM’s (−0.73 kcal/mol), their Tm predictions were nearly the same (AI^2^BMD: 359.61 ± 0.14 K, MM: 359.60 ± 0.13 K) and comparable with the experimental Tm of >372 K^23^. α3D is an artificial protein primarily composed of α-helices^24^. AI^2^BMD’s ΔG estimation (−0.098 kcal/mol) still outperforms MM’s (1.33 kcal/mol). Both the melting temperature of 369.67 ± 0.06 K and 366.94 ± 0.26 K made by AI^2^BMD and MM respectively were comparable to the experimental value (>363 K)^25^. λ-repressor is the largest protein in the evaluation and is dominated by α-helices^26^. AI^2^BMD and MM both deviated from 0 in ΔG estimation (AI^2^BMD: 0.79 kcal/mol, MM: 1.09 kcal/mol), but AI^2^BMD is closer. Their Tm estimations were nearly the same (AI^2^BMD: 349.55 ± 0.21 K, MM: 349.48 ± 0.21 K and closer to the experimental value (347 K)^26^.

For alpha and beta proteins, BBA features a folded structure comprising a α-helix and a β-hairpin^27^. Lacking a known melting temperature, simulations were conducted at 325K. AI^2^BMD’s ΔG calculation (0.057 kcal/mol) and Tm estimation (323.94 ± 0.22 K) were similar with MM’s results (1.22 kcal/mol and 322.34 ± 0.31 K, respectively). Protein G consists of a α-helix and a four-fold β-sheet group^28^. AI^2^BMD’s ΔG prediction (0.14 kcal/mol) is closer to 0 than MM’s (0.74 kcal/mol). In the absence of experimental Tm, AI^2^BMD’s estimated Tm (349.49 ± 0.12 K) demonstrates a 3 K improvement over MM’s (346.49 ± 0.66 K) concerning the simulation temperature of 350 K. Given the free energy and melting temperature evaluations on diverse proteins, AI^2^BMD exhibits well estimations on conformational ensembles, leading to reasonable predictions of protein folding thermodynamics and better alignments for several proteins with experiment data on Tm.

## DISCUSSION

Simulating biomolecular dynamics with *ab initio* accuracy is a long-standing challenge since it is difficult to achieve accuracy, efficiency, and generalization ability at the same time^4,29^. We have developed a generalizable simulation system to all proteins, AI^2^BMD for *ab initio* protein simulation that offers improvements over MM in energy/force calculation accuracy and kinetics/thermodynamic properties estimation. The generalizability of AI^2^BMD across different protein systems and its robustness showcase its potential for broader applications in protein research.

The generalization ability of AI^2^BMD originates from the fundamental principle that most proteins are composed of common kinds of amino acids. This understanding allows AI^2^BMD to serve as a versatile and adaptable algorithm that can be applied to a diverse range of proteins. By incorporating this knowledge into the AI^2^BMD, it can serve for various conformations of a protein and accommodate proteins of different sizes and compositions, enabling researchers to explore and investigate the complex world of proteins with greater confidence and precision.

In the realm of protein dynamics, the *ab initio* accuracy provided by AI^2^BMD has led to the discovery of new insights into the movements and interactions of proteins, gaining a more detailed understanding of the underlying biomechanisms. One notable observation from AI^2^BMD simulation is the increased flexibility in protein movements, which was also observed in MLFF’s simulations^10^. The ability to accurately model the flexibility of proteins is crucial in understanding their functions^30-32^. By incorporating such details into protein simulations, AI^2^BMD can provide a more comprehensive and accurate picture of protein behavior^33^.

While AI^2^BMD boasts a much faster computational speed than DFT methods, it still lags classical MD simulations in terms of efficiency. To bridge this gap, several strategies can be employed in future study. On the one hand, developing a model for all protein units would enable batch inference of energy and force calculations, streamlining the overall process. On the other hand, implementing additional engineering optimization could lead to substantial improvements in the efficiency of AI^2^BMD. Furthermore, applying AI^2^BMD for a broader range of systems, including lipids, nucleotides, nano materials, and solute-solvent interfaces will broaden the scope of AI^2^BMD’s applicability to more complex biomolecular systems to unlock new insights into the intricate world of biomolecular systems^34,35^, paving the way for more accurate and efficient simulations in a variety of contexts, such as drug discovery, protein design and enzyme engineering.

## Author contributions

T.W. led, conceived and designed the study and is the lead contact. T. W., X. H. and M. L. designed the fragmentation approach. X. H. and M. L. built the dataset. Y. W. and S. L. trained machine learning force fields. T. W., X. H., M. L., Y. W. designed the simulation system. S. L, and Z. W. contributed to the simulation system. X. H, M. L., Y. W., S. L. carried out experiments on energy and force calculation for proteins. X. H., M. L. and T. W. carried out kinetics and thermodynamics analysis. B. S. contributed to methodology. T. W., X. H., M. L. and Y. W. wrote the original manuscript. B. S., Z. W., S. L. and T. L. contributed to the manuscript revision. All authors read the final manuscript.

## Acknowledgments

We thank Prof. Haipeng Gong at Tsinghua University and Dr. Frank Noe at Microsoft Research for the constructive discussions.

## Notes

### Competing Interest Statement

The authors have declared no competing interest.

